# SCancerRNA: Expression at the Single Cell Level and Interaction Resource of Non-coding RNA Biomarkers for Cancers

**DOI:** 10.1101/2023.09.26.559661

**Authors:** Hongzhe Guo, Liyuan Zhang, Xinran Cui, Liang Cheng, Tianyi Zhao, Yadong Wang

**Affiliations:** Department of Computer Science, Harbin Institute of Technology, Harbin 150001, China; Molecular Biology Centre, Harbin Medical University, Harbin 150001, China; School of Medicine and Health, Harbin Institute of Technology, Harbin 150001, China

**Keywords:** Non-coding RNA, Biomarker, Cancer, Single cell, Interaction network

## Abstract

Non-coding RNAs (ncRNAs) participate in multiple biological processes associated with cancer as tumor suppressors or oncogenic drivers. Due to their high stability in plasma, urine, and many other fluids, ncRNAs have the potential to serve as key biomarkers for early diagnosis and screening of cancers. During cancer progression, tumor heterogeneity plays a crucial role, and it is particularly important to understand the gene expression patterns of individual cells. With the development of single-cell RNA sequencing (scRNA-seq) technologies, uncovering gene expression in different cell types for human cancers has become feasible by profiling transcriptomes at the cellular level. However, a well-organized and comprehensive online resource that provides access to the expression of genes corresponding to ncRNA biomarkers in different cell types at the single cell level is not available yet. Therefore, we developed the SCancerRNA database to summarize experimentally supported data on long ncRNA (lncRNA), microRNA (miRNA), piwi-interacting RNA (piRNA), small nucleolar RNA (snoRNA), and circular RNA (circRNA) biomarkers, as well as data on their differential expression at the cellular level. Furthermore, we collected biological functions and clinical applications of biomarkers to facilitate the application of ncRNA biomarkers to cancer diagnosis, as well as monitoring of progression and targeted therapies. SCancerRNA also allows users to explore interaction networks of different types of ncRNAs, and build computational models in the future. SCancerRNA is freely accessible at http://www.scancerrna.com/BioMarker.

## Introduction

Despite major advances in precision medicine, cancer continues to be a leading cause of mortality. RNA delivers genetic information and regulates crucial cell processes [1], which can act as a new tool to investigate cancers. Due to their aberrant expression under pathological conditions and ability to circulate in body fluids and maintain paracrine functions [2], non-coding RNAs (ncRNAs) have attracted more and more attention on their potential as cancer biomarkers in clinical trials [3].

Long ncRNAs (lncRNAs) and piwi-interacting RNAs (piRNAs) can serve as important regulators in cancer development [4, 5]. *SCHLAP1* and piRNA-823 can be used as noninvasive biomarkers associated with metastatic progression in prostate cancer and colorectal cancer [6, 7], respectively. Circular RNAs (circRNAs) act as a natural ‘sponge’ by binding and downregulating microRNAs (miRNAs) to influence the cancer disease process [8]. Hsa_circ_0107593 inhibits the proliferation and invasion of cervical cancer cells by sponging three miRNAs [9]. miRNAs are also key regulators of cancer by controlling expression of their target mRNAs to facilitate tumor growth, invasion, and immune evasion [10, 11]. The expression of serum exosomal miR-1226-3p was negatively correlated with the invasion or metastases of pancreatic ductal adenocarcinoma [12]. Small nucleolar RNAs (snoRNAs) are differentially expressed between cancer stages and metastases and can actively modify disease progression [13]. For instance, snoRNAs can function as oncogenes in lung cancer [14] and colorectal cancer [15] to promote abnormal cell growth. Given their vital role in diverse cellular processes, these five types of ncRNA biomarkers have been widely investigated in the diagnosis, prognosis, and personalized therapy aspects of cancers [16].

Accumulating evidence shows that cancer is a highly diverse and heterogeneous disease with distinct histological features, pathologies, and clinical outcome [17]. Abnormal gene expression in different cell types can cause cancer or affect cancer progression. Single-cell RNA sequencing could serve as a powerful tool to detect ncRNA expression and to identify differences between cell subpopulations in human cancers [18, 19]. Due to these characteristics, we can study the genes corresponding to ncRNA biomarkers at the single-cell level. The genes corresponding to biomarkers are the parts of DNA sequences that are transcribed into non-coding RNAs, i.e., non-coding RNA genes. These genes are able to correlated with the non-coding RNA biomarkers. Therefore, a complete and comprehensive ncRNA biomarker database must contain differential expression data at the single-cell level to adequately collect functional annotation information on biomarkers for better use in treatment.

Non-coding RNAs can affect the complexity of progression through their biological functions correlated with cancers [20, 21]. Investigating biological functions and clinical applications of ncRNA brings insight on exploring and utilizing ncRNA biomarkers more comprehensively, which is conducive to the study of cancer occurrence, development and treatment. In addition, an increasing number of machine learning-based models utilize many different RNA types rather than just a single type for disease prediction [22]. Networks in which ncRNAs are involved can influence some molecular targets to drive specific cellular biological responses [23]. Thus, it is crucial to obtain information on the interactions between multiple types of ncRNAs to construct interaction networks for cancer prognosis.

With the development of genomic profiling technologies, several databases based on ncRNA have emerged. Zuo et al. [24] have developed BBCancer to provide expression data on blood-based biomarkers for 15 cancers, but the cancer types are not sufficiently comprehensive, and the sources of biomarkers are relatively singular. Lnc2Cancer [25] documents the information of lncRNA biomarkers comprehensively, but the types of biomarkers are relatively simple. miRandola [26] provides data on miRNAs, lncRNAs, and circRNAs as noninvasive biomarkers but lacks annotation of biological functions and clinical applications. HMDD documents detailed and comprehensive annotations to the human miRNA-disease association data and provides the interaction network, but other types of ncRNA were not involved [27].

Although they are all widely used databases, most of them either focus on one kind of specific ncRNA or lack of annotations and interactions. In addition to the problems mentioned above, none of the existing ncRNA databases provide information on ncRNA genes corresponding to ncRNA biomarkers at the single cell level.

To address the above gaps, we developed SCancerRNA, a manually curated and comprehensive database of five types of ncRNA biomarkers in human cancers, which contains lncRNAs, miRNAs, circRNAs, piRNAs and snoRNAs. SCancerRNA contains differential expression data on genes corresponding to ncRNA biomarkers at the single cell level. In order to better understand the molecular mechanisms underlying the regulation and function of biomarkers in cancer progression and treatment, SCancerRNA provides experimentally supported biological functions (cell proliferation, growth, apoptosis, autophagy and epithelial mesenchymal transformation) and clinical applications (migration, metastasis, circulation, survival and recurrence). To further build models for investigating prognostic prediction and diagnosis of cancers, SCancerRNA provides interaction networks among different types of ncRNAs. SCancerRNA will serve as a comprehensive resource for users to explore the comprehensive collection of ncRNA biomarker types, functional annotations of biomarkers, expression of genes corresponding to biomarkers in different cell types and automatic generation of biomarker interaction networks.

## Data collection and processing

### Data sources

The foundational information on cancer-related ncRNA biomarkers was manually extracted from publications. First, we queried the PubMed database [28] by searching the keywords “non-coding RNA biomarker” and “cancer” to collect five types of cancer-related ncRNA biomarkers, including lncRNAs, miRNAs, circRNAs, piRNAs and snoRNAs. Multiple pieces of information on each ncRNA biomarker were collected from the corresponding literature, such as the type of RNA, synonym of RNA, related cancer type, value (diagnosis, prognosis and therapy), identifier for the source publication and experimental methods (microarray, qRT–PCR, and Western blot, etc.). Subsequently, the reference sequences of lncRNAs, miRNAs, circRNAs, snoRNAs and piRNAs were directly obtained from databases dedicated to each type of RNA. We also classified biomarker-related cancers by tissues, and the 22 tissues corresponding to 219 human cancer subtypes are listed in SCancerRNA.

### Annotation of biomarkers

To comprehensively understand the ncRNA roles of biomarkers, the biological functions and clinical applications corresponding to every biomarker were collected directly from the public literature. According to biological experiments, we divided the biological functions of ncRNAs into cell proliferation, cell growth, apoptosis, autophagy, and epithelial-mesenchymal transition (EMT). The clinical applications of ncRNAs were divided into circulation, survival, recurrence, migration and metastasis.

### Expression of genes based on scRNA-seq

The differential expression information on genes corresponding to ncRNA biomarkers was obtained from published papers. We searched the PubMed database for keywords such as ‘single-cell sequencing cancer’, ‘single-cell sequencing tumor’ and ‘scRNA-seq cancer’. We downloaded the single-cell sequencing literature and the corresponding datasets to collect the genes from different cell types, the value of the average log2-fold-change and the adjusted p value of differential gene expression analysis in different cell types of cancer. In this step, we selected the cell-type-specific genes corresponding to our collection of ncRNA biomarkers. In addition, we manually collected the sequencing platform for each dataset and the steps of quality control during single-cell RNA sequencing.

### Interactions of ncRNAs

Data on regulatory interactions between five types of ncRNAs were obtained from the NPInter [29] database. Since we collected a large number of lncRNAs from existing articles, the number of lncRNA□lncRNA interactions was significantly higher than that of other types of ncRNA interactions. We used the interactions between different types of ncRNAs to automatically generate interaction networks for any one or a group of biomarkers. The whole data collection process is shown in **Figure 1**.

**Figure 1.**
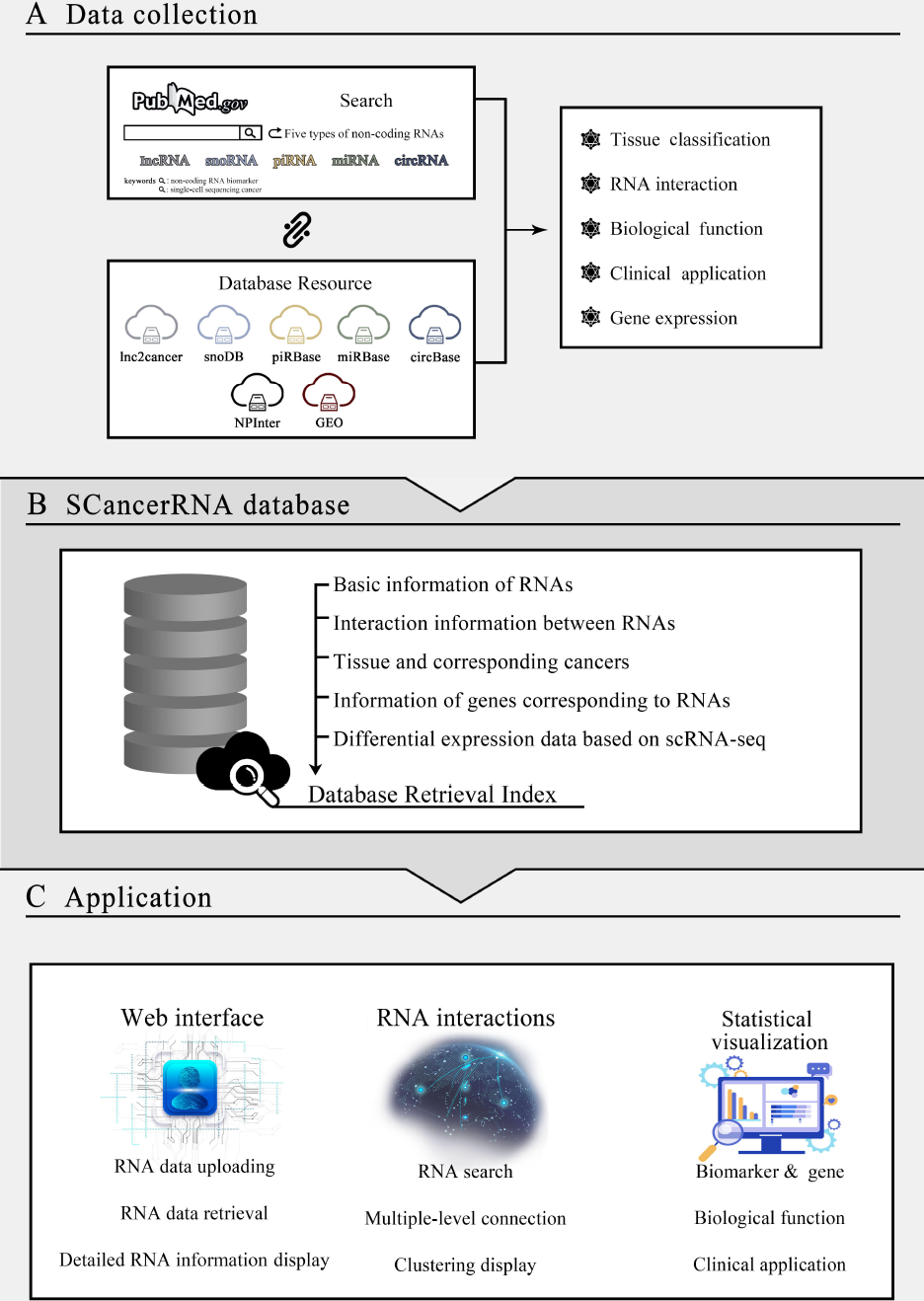
Flowchart of SCancerRNA database construction. **A**. Data collection and processing. **B**. Construction of the SCancerRNA database and retrieval index. **C**. Application of the SCancerRNA database, including the web interface, RNA interactions, and statistical visualization.

## Data content and usage

### Overview of SCancerRNA

The current version of SCancerRNA records 13,381 entries of ncRNA biomarkers by manually curating 7,813 published studies, corresponding to 219 human cancer subtypes. In addition, SCancerRNA documents the expression data of a total of 1,897 genes corresponding to 2,018 ncRNA biomarkers for 19 cancer types at the cellular level by collating 28 single-cell RNA-sequencing datasets from the published literature.

SCancerRNA provides a user-friendly interface that contains seven functions: browse, search, single cell, statistics, download, submit and help. Users are able to obtain five types of ncRNA biomarkers of specific cancers, browse the biological functions and clinical applications of various ncRNA biomarkers, determine the expression of ncRNA genes corresponding to biomarkers in different cell types and explore the automatically generated ncRNA interaction networks (**Figure 2**).

**Figure 2.**
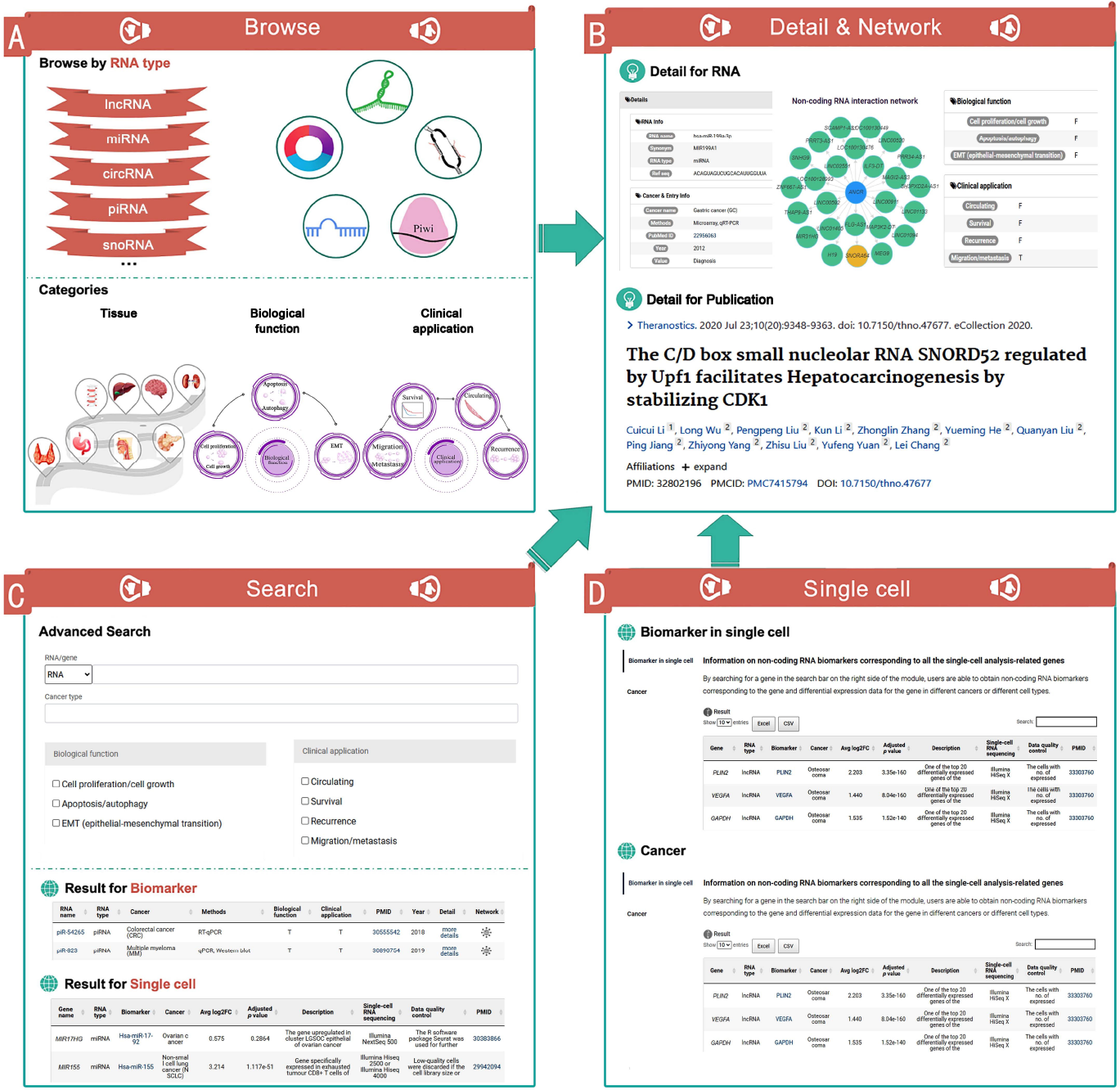
Schematic of the interface in SCancerRNA. **A**. The interface of the browse module with categories of interest. **B**. Modules for detailed information and the interaction network of different ncRNAs. **C**. Search module and interface for displaying search results. **D**. The interface of the single-cell module, ‘Biomarker in single cell’ page and ‘Cancer’ page.

### Browse

SCancerRNA categorized the ncRNA biomarkers by ‘RNA type’, ‘Biological function’, ‘Clinical application’ and ‘Tissue type’. On the Browse page, users can browse SCancerRNA by clicking on diagrams related to the categories listed above, and the matched entries with biomarkers and the genes are returned (Figure 2A). Each entry for the biomarker result includes the name of the ncRNA biomarker, the RNA type of the biomarker, the type of cancer, the testing methods of the ncRNA biomarker, the biological function, the clinical application, the identifier for the source publication, the published year of the reference literature, details (RNA Info, Cancer & Entry Info, Biological Function and Clinical Application) and the interaction network of different types of ncRNA biomarkers.

By clicking on the network link of the queried biomarker, the visualization page (Figure 2B) of the interaction network will pop up, where users can also obtain information on other ncRNAs. If a ncRNA that interacts with the queried ncRNA biomarker is also recorded as a biomarker in SCancerRNA, then clicking on the link of that ncRNA can allow users to obtain the data on the ncRNA in SCancerRNA. Users are also allowed to acquire more detailed information in the original literature by clicking the ‘PMID’ link. Furthermore, each entry of the single-cell result contains 10 sections that describe the relationship among the gene, the corresponding biomarker and the cancer. The 10 sections include the name of the gene, the corresponding biomarker, the RNA type of the biomarker, the type of cancer, the value of the average log2-fold-change, the adjusted p value, the description of the gene in the single-cell differential expression analysis, the platform of scRNA-seq, the steps of data quality control during scRNA-seq and the identifier of the source publication.

### Search

On the ‘Search’ page (Figure 2C), users are allowed to search biomarkers by RNA name and gene name, cancer type or both. Additionally, fuzzy keyword searching functions were provided to better enhance user’s operating experience by returning the closest possible matching records. In this module, users can also select the biological functions and clinical applications of interest to filter the search results for biomarkers. For example, users can narrow the search results by clicking the biological function of apoptosis/autophagy. The format of the search result is the same as that of the browsing result, including the biomarker and the single-cell analysis results.

### Single cell

SCancerRNA provides two modules (Figure 2D) on the ‘single cell’ page, which allows users to easily access biomarkers associated with genes of interest and to discover single-cell expression data associated with specific cancers. In the ‘biomarker in single cell’ module, users are allowed to explore all the ncRNA biomarkers corresponding to the genes related to single-cell differential expression analysis. By searching for a gene in the search bar on the right side of the module, users are able to obtain the differential expression data for the gene in different cancers among different cell types and the SCancerRNA link of the corresponding biomarkers. Furthermore, users can get a detailed understanding of the data quality of each scRNA-seq dataset by checking the data in both ‘Single-cell RNA sequencing’ and ‘Data quality control’ columns in each entry.

In the ‘cancer’ module, users are able to select a cancer type in the cancer drop-down bar on the right to obtain differential expression data for genes associated with the selected cancer at the single-cell level.

### Statistics

The visualization of detailed statistics of SCancerRNA is provided in the “Statistics” function (**Figure 3**). The number of entries corresponding to the five types of ncRNA biomarkers (Figure 3A) is shown in SCancerRNA. The bar chart (Figure 3B and Supplementary Table 1) shows the distribution of biomarker entries of five types of ncRNAs in various human tissues. We also selected several representative ncRNAs (Figure 3C) that could serve as biomarkers for a larger variety of cancers. In addition, the percentage of biological function (cell proliferation, growth, apoptosis, autophagy and epithelial mesenchymal transformation) and clinical application (migration, metastasis, circulation, survival and recurrence) entries (Figure 3D and Supplementary Table 2) for each type of ncRNA biomarker are shown on the statistical page. SCancerRNA also demonstrates the number of genes based on single-cell RNA sequencing in different cancers and the number of their corresponding ncRNA biomarkers (Figure 3E).

**Figure 3.**
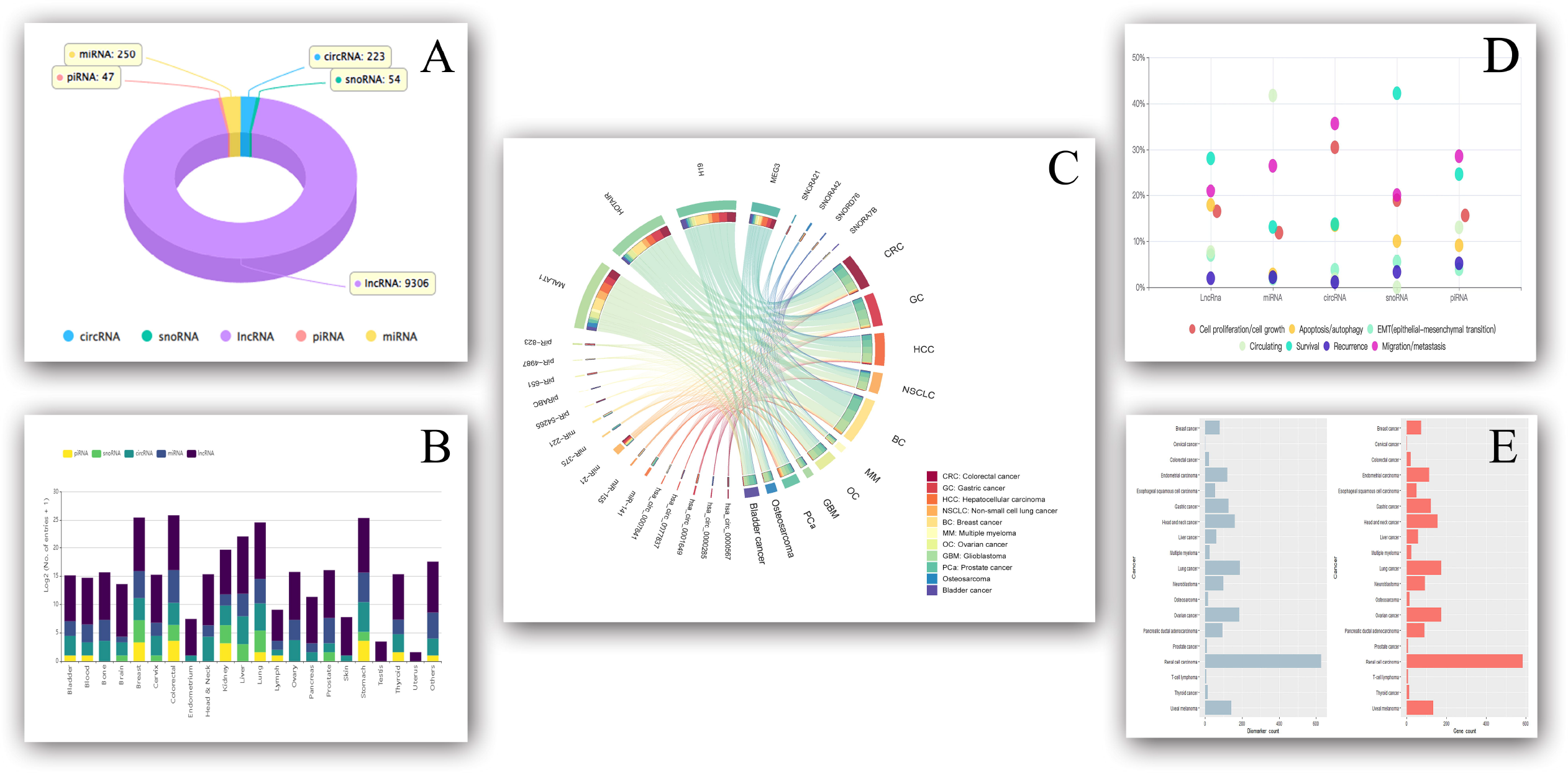
Statistical information. **A**. Data distribution of biomarker entries. **B**. Data distribution of biomarker entries in human tissues. The values of the entries in the chart were log2-transformed after adding a value of one. **C**. Top 5 ncRNA biomarkers of each RNA type with the largest number of corresponding cancer types. **D**. Data distribution of entries on the biological functions and clinical applications of biomarkers. **E**. Data distribution of biomarkers and corresponding genes in cancers

### Download, Submit and Help functions

All the data from the SCancerRNA database can be accessed on the ‘Download’ page, and researchers can upload experimentally supported data on cancer-related biomarkers via the ‘Submit’ function. For new users, the instructions on the ‘Help’ page will make it easier and convenient to explore database.

On the ‘download’ page, seven files are available for users to download. The first five files correspond to detailed entries for five types of ncRNA biomarkers for human cancers. The sixth file is the data on the interactions between different types of ncRNAs, which mainly includes the RNA name, type and interaction score of each pair of interacting ncRNAs. The last file comprises the expression of genes corresponding to biomarkers at the single-cell level, including the value of the average log2-fold-change, adjusted p value, description of the gene in the single-cell analysis, PMID for literature, cancer type, corresponding RNA name and RNA type.

On the ‘Submit’ page, users can submit the novel entry of these five types of ncRNA biomarkers or noninteraction between metabolites and proteins. The RNA name and type of biomarker, corresponding cancer type, methods, reference PMID of the publication and year of publication are needed. The corresponding gene and the Ensembl ID are optional. Users can also select the biological functions and clinical applications of each biomarker to provide more detailed and comprehensive information for SCancerRNA.

On the ‘Help’ page, we provide users with detailed guidance in the form of images or text. Users can get help from this page according to their interests and needs in order to explore SCancerRNA more efficiently.

## Discussion and perspectives

Non-coding RNAs have arisen as key players in the initiation and progression of various cancers, such as modulation of cell proliferation, apoptosis and the cell cycle. With the development of next-generation sequencing, ncRNA biomarkers have been gradually discovered, allowing us to more fully reveal their functions and mechanism of action in various cancers. SCancerRNA is a web-accessible and comprehensive resource that focuses on ncRNA biomarkers and the expression of corresponding genes in multiple human cancers at the single-cell level. It can be freely accessed through a user-friendly web interface: http://www.scancerrna.com/BioMarker.

Compared with other existing biomarker databases, SCancerRNA has the following advantages: (i) it offers multiple types of ncRNA biomarkers for over 200 human cancers, including lncRNAs, miRNAs, circRNAs, piRNAs and snoRNAs; (ii) it contains the differential expression data on genes corresponding to biomarkers at the cell level, which is convenient for users to explore the aberrant expression of biomarkers in human cancers; (iii) it not only provides the experimentally supported biological functions of each ncRNA biomarker but also the clinical applications, adding new dimensions to the understanding of cancer progression and therapy; (iv) it allows researchers to explore the ncRNA interaction network, which facilitates the construction of more accurate computational models of cancer diagnosis and cancer prognosis. Due to rapid advances in molecular biology, we will continue to collect more types of ncRNAs to the database, focusing on their functions and their corresponding non-coding RNA genes differentially expressed among different cell types in cancer. In addition, we hope to construct a more comprehensive ncRNA interaction network to further help researchers discover associations between ncRNAs and cancers.

In recent years, studies on ncRNA biomarkers have been a hotspot in the scientific community. We have collected the experimentally supported entries of ncRNA biomarkers and information on the differential analysis of genes corresponding to the ncRNA biomarkers at the single-cell level. Such information is valuable for gaining further insights into the roles of ncRNA biomarkers in human cancers. Notably, we have also organized the interaction entries and generated the interaction networks of different types of ncRNAs to further construct computational models for cancer prediction. We believe that SCancerRNA will be of particular benefit to the life science community due to the comprehensive collection of ncRNA biomarker types, functional annotations of biomarkers, expression of genes corresponding to biomarkers in different cell types and automatic generation of biomarker interaction networks.

## Data availability

SCancerRNA is free available at http://www.scancerrna.com/BioMarker.

## CRediT authorship statement

**Hongzhe Guo:** Conceptualization, Methodology, Formal analysis, Software, Visualization, Writing – original draft, Funding acquisition. **Liyuan Zhang:** Validation, Investigation, Writing - Review & Editing. **Xinran Cui:** Data curation, Investigation. **Liang Chen:** Methodology, Resources. **Tianyi Zhao:** Conceptualization, Supervision, Funding acquisition. **Yadong Wang:** Conceptualization, Resources, Supervision, Project administration. All authors read and approved the final manuscript.

## Acknowledgments

This work has been supported by the National Natural Science Foundation of China (Grant No. 62102116 and 62202125) and the Interdisciplinary Research Foundation of Harbin Institute of Technology.

## Figure legends

**Supplementary Table 1 Tissue distribution of ncRNA biomarkers collected in SCancerRNA**

**Supplementary Table 2 Experimentally supported biological functions and clinical applications of the five types of noncoding RNA biomarkers**.

